# Experimental evidence that animal societies vary in size due to sex differences in cooperation

**DOI:** 10.1101/2022.03.01.482465

**Authors:** Julian Melgar, Mads F. Schou, Maud Bonato, Zanell Brand, Anel Engelbrecht, Schalk W. P. Cloete, Charlie K. Cornwallis

## Abstract

Cooperative breeding societies allow the costs of reproduction to be shared. However, as groups become larger, such benefits often decrease and competition increases. This is predicted to select for an optimal group size, yet variation in groups is a ubiquitous feature of cooperative breeding animals. Here we experimentally established groups (n_groups_=96) of cooperative breeding ostriches, *Struthio camelus*, with different numbers of males and females and manipulated the potential for cooperation over incubation. There was a clear optimal group size for males. Their reproductive success was maximized in groups with four or more females and no other males, irrespective of cooperation over incubation. Conversely, female reproductive success was strongly dependent on the benefits of cooperating over incubation, being maximized in groups with either many males or many females. In intermediate sized groups, both male and female reproductive success was reduced by sexual conflict over the timing of mating and incubation. Our experiments show that sex differences in the opposing forces of cooperation and competition can explain why variation in cooperative groups is widespread.

## Introduction

Group size is a key feature influencing the social organization of cooperative breeding animals (Alexander, 1974; Bourke, 1999; Clutton-Brock, 2021; Kappeler, 2019; Koenig and Dickinson, 2016; Rubenstein and Abbot, 2017; Taborsky et al., 2021). In large groups there are greater opportunities for cooperation that can increase individual reproductive success, for example, by spreading the burden of offspring care among group members (Alexander, 1974; Koenig and Dickinson, 2016; Rubenstein and Abbot, 2017; Taborsky et al., 2021). However, as groups increase in size, higher competition and lower marginal benefits of cooperation can reduce reproductive success, especially in cooperative breeding systems where all group members attempt to breed (Clutton-Brock, 2021; Krause et al., 2002; Powers and Lehmann, 2017; Riehl, 2011; Vehrencamp, 1983). Consequently, it is predicted that the counteracting forces of cooperation and competition will lead to the evolution of optimal group sizes (Fig. 1. Krause et al., 2002; Markham et al., 2015; Powers and Lehmann, 2017; Pulliam and Caraco, 1984).

**Fig. 1:**
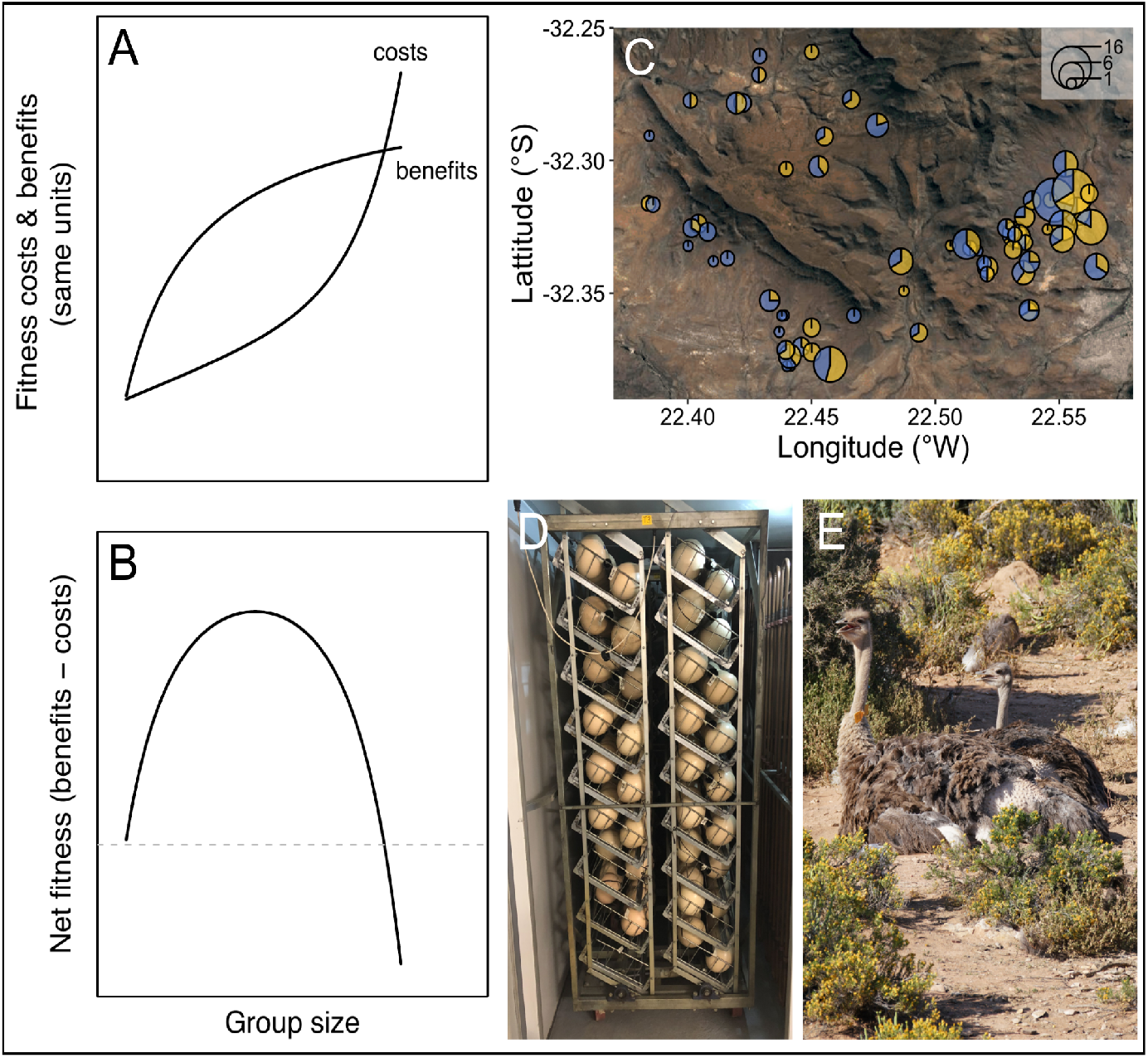
Is there an optimal group size for cooperative breeding ostriches? Theoretically, accelerating costs of competition and diminishing benefits of cooperation are expected to result in a single optimal group size (A-B, modified from Krause et al., 2002). (C) Groups in natural populations are, however, highly variable in size: A map of the Karoo National Park with different groups of ostriches plotted. The size of the circles indicate the number of individuals and the blue and yellow segments indicate the proportion of males and females respectively. To understand natural variation in group size, experiments manipulating the number of males and females in groups, and the benefits of cooperation are required. Opportunities for cooperation over incubation were manipulated at the experimental study site by collecting and artificially incubating eggs (D). Patterns of reproductive success in relation to group size when cooperation was restricted were compared to situations where opportunities for cooperative incubation, such as among these females (E), were allowed.

Cooperative breeding groups are, however, extremely variable, differing in their numbers of males and females (Alexander, 1974; Clutton-Brock, 2021; Koenig and Dickinson, 2016; Lott, 1991; Lukas and Clutton-Brock, 2018; Rubenstein and Abbot, 2017; Rudolph et al., 2019). One explanation is that optimal group sizes are difficult to observe due to the complexity of natural systems. For example, fluctuating ecological conditions may shift the optimal size of groups over time and space (Koenig, 1981; Yip et al., 2008; Zöttl et al., 2013).

Alternatively, assumptions about the way the costs and benefits change with group size may need revising (Giraldeau and Gillis, 1985; Krause et al., 2002; Pulliam and Caraco, 1984; Sibly, 1983; Yip et al., 2008). Empirical estimates of the costs and benefits of group size are typically inferred from measuring the outcome of breeding events from natural groups (Alexander, 1974; Koenig and Dickinson, 2016; Rubenstein and Abbot, 2017; Taborsky et al., 2021). However, it is difficult to use such data to quantify the benefits and costs to individuals of being in different sized groups. This is because the effects of competition over mating cannot be separated from the effects of cooperation over offspring care, and estimates of reproductive success can be biased by individual differences (Cockburn et al., 2008; Dickinson and Hatchwell, 2004; Downing et al., 2020; Schoepf and Schradin, 2012). For example, the benefits of being in large groups can be overestimated if individuals with high reproductive success are more likely to be in large groups. Males and females can also differ in their optimal group sizes due to divergent reproductive interests (Davies, 1989; Lessells, 2012; Trivers, 1972; Wong et al., 2012), but understanding how these effects combine to determine group size variation is difficult without experimental manipulations.

Experiments are therefore needed to control for ecological conditions, partition individual differences in reproductive success from group level effects, and separate how competition and cooperation change with group size. Such experiments are extremely challenging to conduct, particularly in large vertebrates (Krause et al., 2002). We overcome these challenges by studying cooperative breeding ostriches, *Struthio camelus*, where experimental manipulations are possible. Ostriches breed in groups ranging from pairs to large multi-male, multi-female groups, which have a communal nest where all individuals typically attempt to breed (Fig. 1. Bertram, 1992; Sauer and Sauer, 1966). Both males and females participate in cooperative incubation of eggs for 42 days, representing a major part of parental care (Kimwele and Graves, 2003; Magige et al., 2009). During this period, eggs must be constantly protected exposing adults to risks of heat exhaustion and predation (Bertram, 1992; Magige et al., 2008; Sauer and Sauer, 1966). Using this system, we first quantified group sizes in wild populations, and second experimentally tested how group size variation influences male and female reproductive success (n_groups_ = 96, n_individuals_=273).

At the start of each breeding season, groups were experimentally established by placing individuals in large enclosures in the Klein Karoo, South Africa. Groups consisted of one to three males and one to six females, in accordance with the core range of group sizes seen in the wild (Fig. 1. Table S1). The number of males and females in groups were varied independently, enabling sex differences in optimal group size to be estimated, and interactions between the number of each sex (sex ratio) to be accounted for. To separate the costs of sexual competition from the benefits of cooperation in groups of different size, we measured reproductive success both with and without opportunities for cooperation over incubation. Opportunities for cooperative incubation were restricted by experimentally removing eggs from their nests and hatching them in artificial incubators (Fig. 1). We collected data on the number of chicks produced over a period of five months where eggs were artificially incubated, and two months where eggs were naturally incubated (data on the number of eggs produced are presented in the supplementary material: Fig. S1, Tables S2-S3). This allowed us to estimate the mean reproductive payoffs for males and females in each group (total number of chicks / number of same sex individuals in the group) with and without opportunities for cooperation over incubation.

## Results

Groups of wild ostriches were highly variable in their sizes. Groups observed in the Karoo National Park consisted of one to twelve individuals of the same sex, most often containing one to six females and one to three males (Fig. 1. Table S1). This is similar to the composition of groups reported in East African populations, showing that local variation in groups is widespread across their geographical distribution and across populations (Bertram, 1992; Kimwele and Graves, 2003; Magige et al., 2009).

### Sexual competition regulates the optimal group size for males

In experimental groups when opportunities for cooperation over incubation were restricted, male reproductive success increased with the number of females and decreased with the number of males (Fig. 2A. Number of females *β* (credible interval: CI) = 0.54 (0.26, 0.79), pMCMC = 0.001. Males 3 versus 1 *β* (CI) = -1.24 (−1.7, -0.86), pMCMC = 0.001. Table S4). Allowing for cooperation over incubation reduced the negative effect of competition on male reproductive success, particularly in groups with fewer females (Fig. 2A-B. Number of females*Number males *β* (CI) = -2.12 (−4.05, -0.12), pMCMC = 0.008. Table S4), and further increased the benefits of extra females for single males (Fig. 2A-B. Single males: number of females, care versus no care *β* (CI) = -1.74 (−4.1, -0.22), pMCMC = 0.008. Table S4). However, this did not change the optimal group size for males: Males produced most offspring when they were in groups on their own with four or more females, regardless of the need of cooperation over incubation. This illustrates that male competition and the availability of females are the predominant forces shaping male reproductive success.

**Fig. 2:**
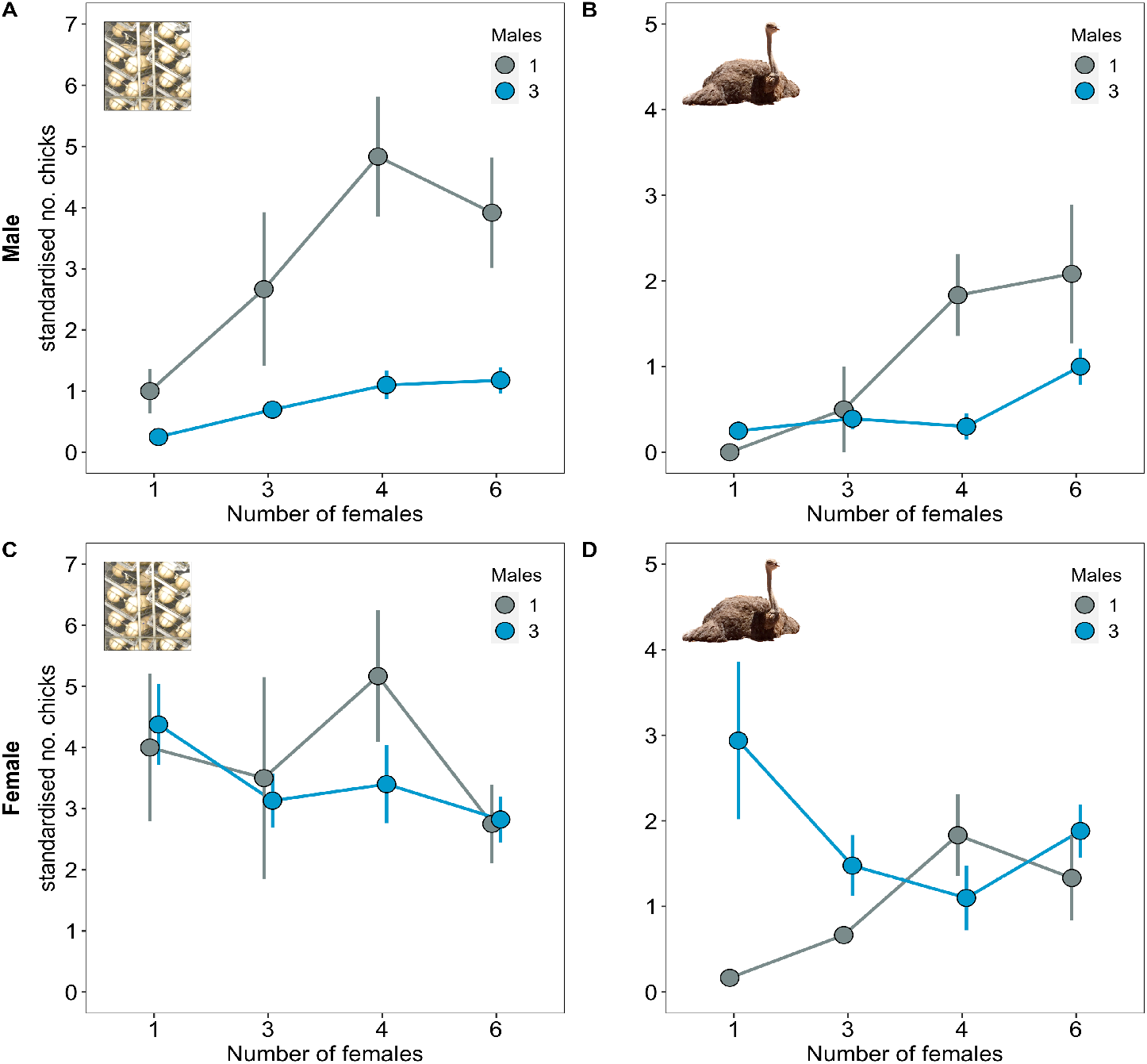
Group size and opportunities for cooperation over incubation influence male and female reproductive success. Reproductive success was measured as the number of chicks produced per individual per clutch for each reproductive stage (no care versus care. See methods for details). (A) The number of chicks males sired decreased with the number of males in the group and increased with the number of females. (B) Opportunities for cooperation over incubation reduced the effects of male competition in groups with few females and magnified the effect of the number of females in groups without male competition (Table S4). (C) When opportunities for cooperative incubation were removed the number of chicks females produced was independent of the number of males and females in groups. (D) When there were opportunities for cooperative incubation, the number of chicks females produced was dependent on both the number of males and females in groups. Means ± SE are plotted.

### Cooperation results in multiple group size optima for females

When the need for incubation was removed, female reproductive success was independent of the number of males and females in groups (Fig. 2C & S1. Males 3 versus 1 *β* (CI) = -0.1 (−0.39, 0.35), pMCMC = 0.856. Number of females *β* (CI) = -0.1 (−0.28, 0.04), pMCMC = 0.13. Table S5). However, when there were opportunities for cooperative incubation, female reproductive success was strongly dependent on both the number of males and the number of females in groups (Fig. 2D. Table S5). In groups with single males, the number of offspring females produced increased linearly with the number of females (Fig. 2D. Number of females *β* (CI) = 0.55 (0.04, 1.74), pMCMC = 0.008. Table S5). In contrast, in groups with three males, females produced more offspring when on their own and in groups with six females (Fig. 2D. Groups with 3 males versus 1 male: number of females^2^ *β* (CI) = 1.01 (0.2, 1.68), pMCMC = 0.004. Table S5). Female reproductive success was therefore maximised across multiple group compositions due to the benefits of cooperating over incubation with other male and female group members (Fig. 2. Table S5).

### Cooperative care in larger groups increases hatching success and lightens workloads

We investigated how cooperation over incubation influenced the group size optima for males and females by observing their behavior, and found that in multi-male and multi-female groups, individuals frequently shared incubation. This led to the total time eggs were protected increasing with the number of males and females in groups (Fig. 3A. Males 3 versus 1 *β* (CI) = 1.27 (0.29, 2.06), pMCMC = 0.006. Number of females *β* (CI) = 0.94 (0.56, 1.34), pMCMC = 0.001. Table S6). In turn, hatching success was higher in groups where nests were protected for longer (Fig. 3B. *β* (CI) = 0.35 (0.03, 0.7), pMCMC = 0.034. Table S7). Individuals did not, however, spend more time incubating in larger groups (Tables S8 & S9). In fact, males spent less time incubating when there were other males, despite nests being protected for longer (Fig. 3C & 3D. Males 3 versus 1 *β* (CI) = -3.07 (−5.28, -0.76), pMCMC = 0.004. Table S8). It therefore appears that the advantage of being in larger groups is explained by cooperation over incubation spreading the load of parental care and increasing hatching success.

**Fig. 3:**
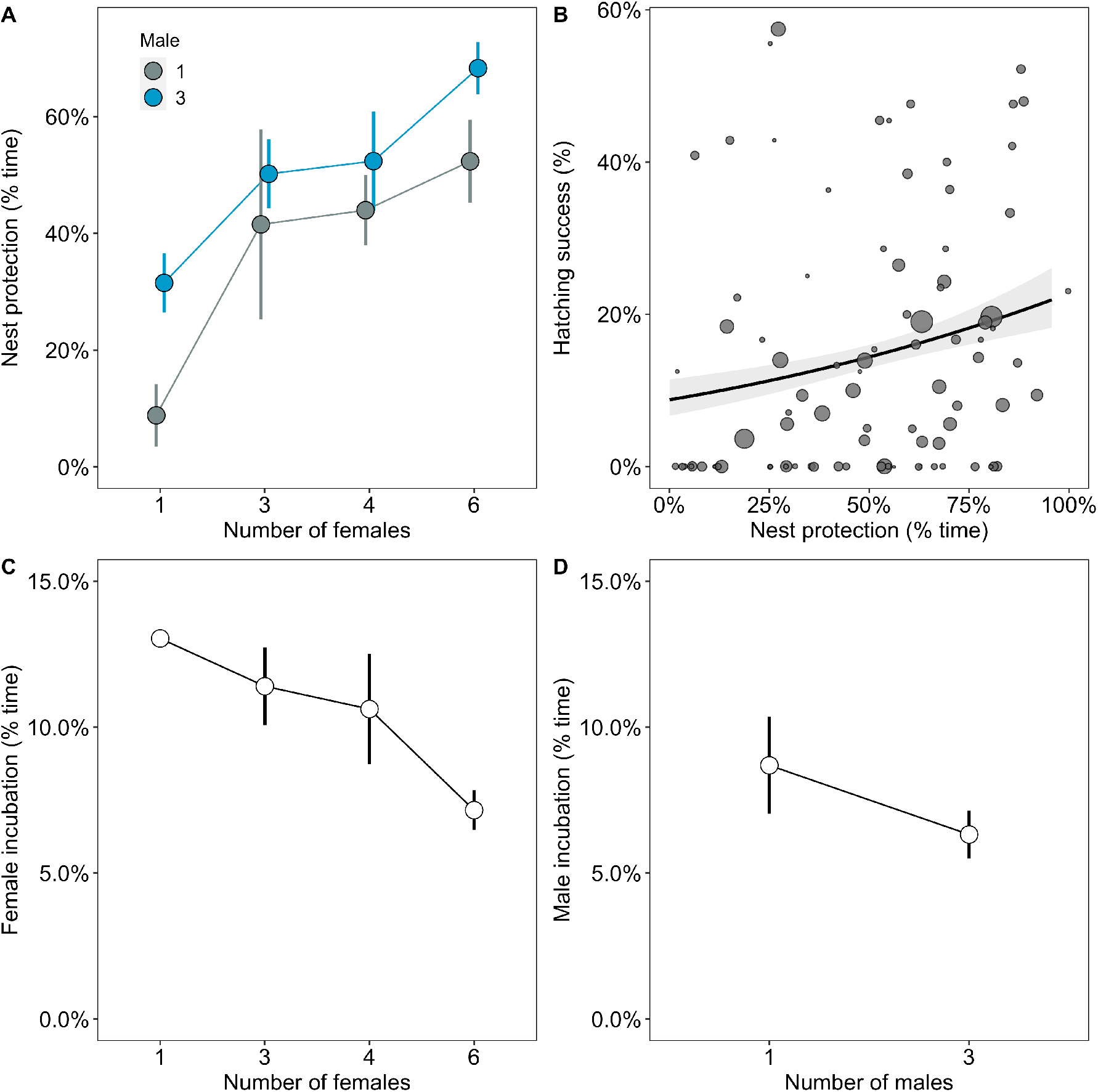
The benefits of cooperative parental care in relation to group size. (A) The amount of time nests were protected was higher in groups with more males and females. (B) Hatching success increased with the amount of time nests were protected. Regression line from a binomial generalized linear model (GLM) with 95% confidence intervals is shown and the size of the points represent the number of eggs laid by groups. (C) The amount of time females spent incubating decreased with the number of females in groups, although not significantly. (D) Males spent less time incubating in groups with three males compared to when they were on their own. Means ± SE are plotted in A, C and D.

### Sexual conflict over the timing of mating and incubation disfavours intermediate group sizes

Limited opportunities for cooperation over incubation may constrain the reproductive success of individuals in small groups (Fig. 2 & 3). However, this does not explain why reproductive success was low in groups of intermediate size. In some groups, males were seen initiating copulations with females that were incubating. This disturbed incubation and resulted in eggs being displaced and broken. We investigated whether such a lack of coordination over mating and incubation explains reductions in the reproductive success of individuals in groups of intermediate size.

The number of interruptions to incubation increased with the number of males in groups (Fig. 4A. Males 3 versus 1 (CI) = 1.5 (0.5, 2.73), pMCMC = 0.002. Table S10). This was most pronounced in groups with intermediate numbers of females (Fig. 4A. Number of females^2^ *β* (CI) = -0.67 (−1.13, -0.16), pMCMC = 0.018. Table S10). In contrast, in groups with the lowest and highest numbers of females, interruptions to incubation were relatively rare (Fig. 4A. Table S10). Interruptions to incubation were associated with a mismatch in the amount of time males and females spent incubating, but only in groups where there was male competition (Fig. 4B. Differences in incubation: 3 males *β* (CI) = -1.43 (−2.36, -0.26), pMCMC = 0.024, 1 male *β* (CI) = 0.18 (−1.4, 1.33), pMCMC = 0.994. Table S11). Interruptions were therefore more frequent in groups with multiple males where females incubated for longer than males, but not where males invested more time than females incubation (Fig. 4B).

**Fig. 4:**
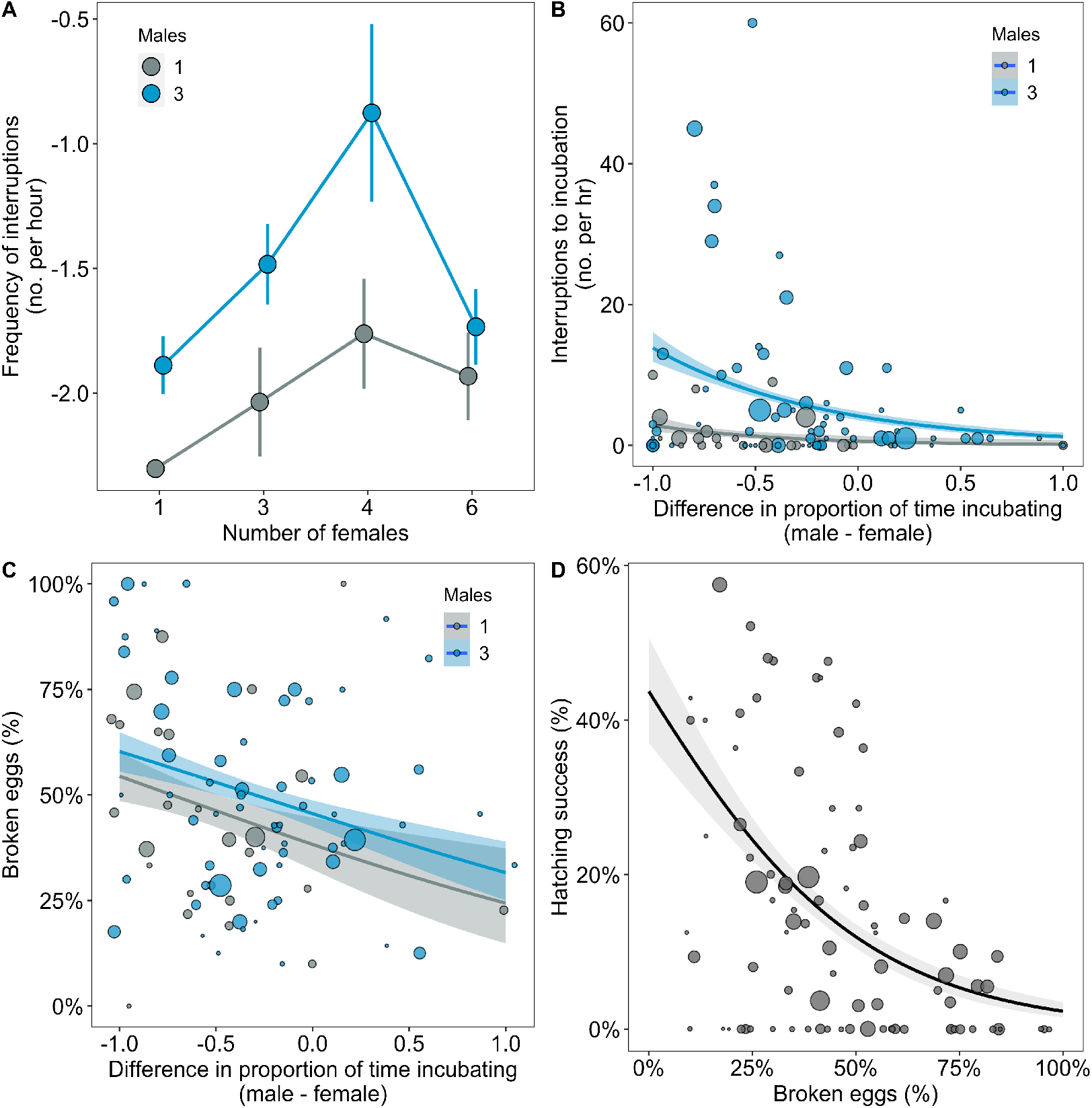
Coordination over reproduction changed with group composition. (A) The frequency of interruptions during incubation increased with the numbers of males in groups, especially when there were intermediate numbers of females. Means ± SE are plotted. (B) Interruptions to incubations were more frequent in groups with three males when females spent more time incubating than males. (C) More eggs were broken in groups with greater disparities in the time males and females spent incubating, which decreased hatching success (D). Regression lines are from GLMs (B = Poisson; C & D = Binomial) with 95% confidence intervals. The size of the points in B, C and D represent the number of eggs laid by groups.

Disparities in incubation between males and females influenced the probability of egg breakages. Eggs were more frequently destroyed when females invested more time in incubation than males (Fig. 4C. Differences in incubation in groups with 3 males: *β* (CI) = -0.64 (−1.23, -0.06), pMCMC = 0.026. Table S12), which reduced hatching success (Fig. 4D. *β* (CI) = -0.82 (−1.2, -0.58), pMCMC = 0.001. Table S13). A lack of coordination between males and females over the timing of mating and incubation therefore appears to explain why more eggs were broken in groups with intermediate numbers of males and females. Conflicts over the timing of reproduction between the sexes have been found to influence reproductive success in other species (Holland and Rice, 1998; Løvlie and Pizzari, 2007). Our results now show such conflicts may also be an important factor shaping the composition of cooperative breeding groups.

## Discussion

Our experiments show that the numbers of males and females in groups have pronounced effects on reproductive success, helping explain the prevalence of group size variation in natural populations. While the change in the costs and benefits of increasing numbers of males and females resulted in a clear optimal group size for males, this was not the case for females. The benefits of cooperative incubation and the costs of sexual conflict lead to female reproductive success being maximised across multiple group sizes. Although the importance of competition and cooperation for group living species has long been recognized (Alexander, 1974; Williams, 1966), our results demonstrate that differences in their relative strengths in males and females can select for different group sizes and compositions.

In species where group members are typically unrelated, ongoing sexual conflict over parental care and parentage is generally thought to explain variation in breeding systems, both within and between species (Davies, 1989; Lessells, 2012; Trivers, 1972; Wong et al., 2012). For example, in dunnocks, *Prunella modularis*, and alpine accentors, *Prunella collaris*, female reproductive success is increased by the number of males providing care, but male reproductive success declines due to more intense sexual competition (Davies et al., 1995; Davies, 1992; Davies, 1985; Hartley and Davies, 1994; Santos and Nakagawa, 2013). Consequently, polyandrous breeding systems are thought to arise where females have the upper hand, and polygynous breeding systems where males gain control (Davies, 1989).

Sexual conflict also appears to influence the types of social groups observed in ostriches. The pursuit of reproductive opportunities by males beyond the point where females engage in parental care appears to lead to a ‘sexual tragedy of the commons,’ causing the demise of groups with intermediate numbers of males and females (Galliard et al., 2005; Hardin, 1968; Rankin et al., 2011). However, sexual conflict appears to play a relatively minor part in explaining overall variation in social groups. Instead, differences in the way the benefits of cooperation and the costs of sexual competition are balanced by males and females appears to generate a broad range of group sizes where the reproductive payoffs are similar.

Our results show that monitoring only the outcome of breeding events makes it difficult to estimate the relative contributions of mate choice, sexual competition and cooperation to selection for group living. For example, without manipulating the need for cooperation it would have been difficult to ascertain if the elevated success of single females in groups with three males was because of differences in mate choice opportunities, or because of the extra parental care provided by males. Given that the reproductive success of single females did not vary with the number of males when opportunities for cooperation were restricted, it appears that female reproductive success is increased by paternal care and not by mate choice opportunities. This highlights the importance of experiments in identifying the sources of selection shaping social organization of cooperative breeding animals.

Under natural conditions, group sizes are likely to depend on a variety factors, such as predator defence and competition over food (Alexander, 1974; Koenig and Dickinson, 2016; Rubenstein and Abbot, 2017; Taborsky et al., 2021). Evidence from mammals, fish, birds and arthropods shows that the size of cooperative groups can vary with ecological conditions (Bertram, 1980; Bourke, 1999; Clutton-Brock, 2021; Koenig, 1981; Yip et al., 2008; Zöttl et al., 2013). However, our results demonstrate that differences in cooperative breeding groups can arise independently of the main factors currently used to explain variation in cooperative breeding groups: ecological conditions, breeder quality and relatedness (see also Casari and Tagliapietra, 2018). This highlights the importance of establishing how social interactions influence selection on sociality when interpreting patterns of social organisation in nature.

## Materials and Methods

### Study population

This study was conducted on two populations. Natural variation in group composition was examined in a wild population of ostriches in Karoo National Park, South Africa (32°19“49.27”S, 22°29’59.99”E) in 2014 (8-9th November) and 2018 (17-19th November). The experiments manipulating group size were conducted on a captive population of ostriches kept at Oudtshoorn Research Farm, South Africa (33° 38’ 21.5”S, 22° 15’ 17.4”E) from 2012 to 2018.

### Natural variation in group composition

Natural variation in the composition of breeding groups (group size, number of males and number of females) was examined using published literature (Bertram, 1992; Kimwele and Graves, 2003; Magige et al., 2009), and directly estimated by conducting transects along the roads of the south eastern part of Karoo National Park. Each transect was carried out 2 to 3 times. Ostriches were typically observed in clearly defined groups, judged by their coordinated movement and close proximity to each other (< ∼100m). In a few instances, individuals were separated by more than 100m. In these situations they were observed until it was clear whether they were part of a group or were moving separately. Whether individuals were sexually mature (immature females = no or very few white wing feathers; immature males = mix of brown and black body plumage) was also recorded. Immature individuals were observed only twice; one group of three males and one female, approximately two years old, and one group of seven individuals, approximately 1 years old and therefore not possible to sex. Fig. 1B includes the two-year-old group, but not the one-year-old group.

### Experimental design

We experimentally manipulated the composition of 97 groups of breeding ostriches involving 280 adult ostriches (127 males and 153 females), over a seven-year period (10-16 groups per year determined by the availability of birds and enclosures). Groups were kept in fenced areas (range: 2400 and 70600 m^2^, median = 4700 m^2^) at Oudtshoorn Research Farm (Cloete et al., 2008) and were randomly allocated one or three males and one to six females. Due to limitations in the number of birds accessible for our experiments, and other experiments being conducted on the same population, not all combinations of male and female group sizes were possible (see Table S16 for full details). All individuals in the Oudtshoorn population were individually identifiable by coloured and numbered neck tags.

In seven groups, an individual was injured or died part way through the season and was replaced by a new individual. In one case the injured individual was the only male in the group and we therefore only included data from the point when the replacement male was introduced (see Table S17). In 12 groups, spread across years and group sizes, it was not possible to replace injured individuals. The number of individuals was consequently reduced (seven groups = six to five, one group = four to three, three groups = three to two. See Table S17 for details of replacements and removals). To avoid creating new group size treatments with low sample sizes, these groups were treated as part of their intended group size treatments in the analyses. One group was excluded from all analyses as the injured individual was the only female in the group. Our final sample size was therefore 96 groups (Table S16).

The breeding season was typically from May to December every year. During the first ∼5 months of the season, eggs were collected to measure reproductive success independently from the effects of incubation behaviour. During the last ∼2 months, eggs were left in nests and incubation behaviour was monitored to examine patterns of reproductive success when individuals had to care for offspring. Reproductive success was measured as the number of eggs and number of chicks produced by groups. During the breeding season ostriches received a balanced ostrich breeder diet (90 to 120 g protein, 7.5 to 10.5 MJ metabolizable energy, 26 g calcium and 6 g phosphorus per kg feed) and ad-libitum water.

### Measuring reproductive success when opportunities for cooperative incubation were removed

To measure reproductive success independently of incubation behaviour, eggs were collected from nests twice a day and artificially incubated. Eggs were marked according to the time of day, date and group of origin, and placed under UV lights for 20 minutes for disinfection. As eggs were incubated in batches starting once a week, eggs were stored prior to incubation for one to six days under conditions known to maintain hatching success (Brand et al., 2008): Eggs were kept on turning trays (two daily 180° rotations) in a cold room (17°C) with relative humidity between 80% and 90%. Hereafter eggs were transferred to artificial incubators set at 36.2°C with a relative humidity of 24%. Eggs in the incubator were automatically turned 60° around their long axis every hour. Eggs were inspected daily for signs of pipping from day 39 of incubation. The incubation period in ostriches is ∼42 days. We were interested in the average reproductive returns for individuals in groups of different size, irrespective of between individual variation in reproductive success within groups. Therefore, we estimated individual reproductive success as the number of eggs and chicks produced by groups, divided by the total number of same sex individuals within groups.

### Measuring reproductive success when there were opportunities for cooperative incubation

Nests were checked daily and new eggs were marked with the date and an egg identification number. The absence and presence of previously laid eggs was recorded to track the fate of each egg. During this period, the incubation behaviour of individuals was monitored by conducting ∼3 hour observations at least three times a week using binoculars (10 × 40) and a telescope (12-36 × 50). The observer sat camouflaged in a ten meter tall observation tower in the middle of the field site. Groups were observed for 60.54±1.38 hours (mean±se) spread over the season. The identity of each incubating individual, as well as the start and end of incubation, were recorded. When incubation was interrupted by other individuals in the group, the time of the interruption and the identity and sex of the interrupting individual was recorded. The consequences of interruptions varied in severity from individuals returning to nests within seconds to individuals ceasing incubation for that observation period. To avoid including minor disturbances in our measure of the number of interruptions, we only included interruption events that resulted in the incubating individual not returning to the nest within one minute.

In the first three years (2012 to 2014), hatching success was measured by allowing groups to naturally incubate eggs to completion. If no eggs were observed hatching in groups after 50 days of incubation, they were removed and checked for developing embryos. From 2015 onwards, changes in legislation to reduce the spread of avian flu meant that contact between adults and chicks had to be minimised. Consequently, eggs were removed from nests just before hatching (∼40 days after the onset of incubation) and placed in artificial incubators to determine hatching success. Individual reproductive success was estimated in the same way as when parental care was removed: the number of eggs and chicks produced by groups divided by the total number of same sex individuals within groups.

### Standardisation of reproductive success

To be able to compare patterns of reproductive success between periods when eggs were artificially and naturally incubated and across different years with slightly different breeding season lengths, we standardised the number of eggs and chicks produced by individuals. First, the number of eggs individuals produced was divided by the number of days reproduction was monitored to get a measure of reproductive output per day. Second, measures of individual daily egg output were divided by the maximum egg output for that sex for that stage of the experiment (no care versus care). This was important because hatching success was much higher when eggs are artificially incubated compared to natural incubation. This gave a maximum egg laying rate per day of one for both stages. However, females can biologically only produce one egg every two days. We therefore divided our standardised measures by two and multiplied by the number of days it typically takes to form a clutch (20 days) in order to give an indication of the eggs laid per clutch. The exact same procedure was used to standardise the number of chicks individuals produced.

### Statistical analyses

#### General approach

Data were analysed in R (R Core Team, 2020) using Bayesian Linear Mixed Models (BLMM) with Markov chain Monte Carlo (MCMC) estimation in the package MCMCglmm v. 2.29(Hadfield, 2010). Default fixed effect priors were used (independent normal priors with zero mean and large variance (10^10^) and for random effects inverse gamma priors were used unless otherwise specified (V = 1, nu = 0.002). Each analysis was run for 3e+06 iterations with a burn-in of 1e+06 and a thinning interval of 2000. Convergence was checked by running models three times and examining the overlap of traces, levels of autocorrelation, and testing with Gelman and Rubin’s convergence diagnostic (potential scale reduction factors <1.1)(Brooks and Gelman, 1998).

Parameter estimates (*β*) for fixed effects were calculated using posterior modes and are reported from models that included all terms of the same order and lower. For example, all main effect estimates are from models where all other main effects are included, all estimates of two-way interactions are from models that included all two-way interactions and main effects, and so forth. Quadratic effects were tested in models including main effects and effects of the same order (other quadratic effects and two-way interactions). The length of time groups were monitored was accounted for by including it as a fixed effect. All continuous explanatory variables were z transformed using the ‘scale’ function in R. Explanatory variables that were proportions were logit transformed using the ‘logit’ function in R and count variables were log transformed. Curvilinear effects of continuous explanatory variables were modelled using the quadratics of the z transformed values computed before running the models.

Fixed effects were considered significant when 95% credible intervals (CIs) did not overlap with 0 and pMCMC were less than 0.05 (pMCMC = proportion of iterations above or below a test value correcting for the finite sample size of posterior samples). By default MCMCglmm reports parameter estimates for fixed factors as differences from the global intercept. This does not allow absolute estimates and 95% CIs for all factor levels to be estimated or custom hypothesis tests of differences between factor levels. Consequently, we removed the global intercept from all models and present absolute estimates for factor levels. Differences between factor levels were estimated by subtracting the posterior samples from one level from the second level and calculating the posterior mode, 95% CI and pMCMC.

Random effects were used to model the non-independence of data arising from multiple data points per individual per group, per enclosure and per year. Random effect estimates presented in tables are from models that included the highest order fixed effect terms. To estimate the magnitude of random effects we calculated the percentage of the total random effect variance explained by each random term on the expected data scale (I2%: (V_i_/V_total_) * 100)(Villemereuil et al., 2016). To obtain estimates of I2 on the expected scale from binomial models the distribution variance for the logit link function was included in the denominator: (V_i_/V_total_ + *π*^3/2^)*100.

#### Specific analyses

##### 1. Testing how group size and the need for parental care influences male and female reproductive success

The effect of group composition on the standardised number of eggs individuals produced was modelled using a BLMM with a Poisson error distribution. The need for parental care (2 level factor: no care vs care), number of males (two level factor: one versus three), the number of females (continuous) and the time groups were monitored (continuous) were entered as fixed effects, and year and enclosure were included as random effects. The effects of group composition on the number of eggs produced per adult bird with and without the need for parental care, were estimated by fitting two-way interactions between care and the number of males and females in groups. Separate models were run for males and females (R code: M1 & M2). The number of chicks males and females produced was modelled in exactly the same way (R code: M3 & M4).

##### 2. Testing how the benefits of cooperative parental care vary with group size

The effect of group composition on the proportion of time groups protected nests was modelled using a BLMM with a binomial error distribution. The response variable was the number of observation minutes birds were sitting on nests versus the number of observation minutes nests were exposed. This accounts for variation across years in observation effort. The number of males and females in groups were included as fixed effects and year and enclosure were random effects (R code: M5). The effect of the proportion of time nests were protected on hatching success was modelled using a BLMM with a binomial error distribution of the number of eggs hatched per individual (total number of chicks produced by groups / number of same sex individuals) versus the number of eggs that did not hatch per individual (total number of eggs that failed to hatch / number of same sex individuals). The proportion of time nests were protected, the number of males and females in groups and the amount of time groups were monitored were included as fixed effects Year and enclosure were included as random effects (R code: M6).

Typically groups only had one active nest, but in a few cases a second and a third nest were occasionally used. The amount of time groups protected their nests was calculated by summing data across all nests (total time nests were protected versus total time nests were exposed). Data were summed across nests to facilitate comparisons with the egg and chick data, which were recorded at the level of the group (e.g. total number of eggs and chicks groups produced by each group), not at the level of each nest. To check if the number of nests groups used influenced the time nests were protected and hatching success, we included the number of nests (continuous) as a fixed effect (R code: M5 & M6). The number of nests did not have a significant effect in any of our analyses (Tables S6 & S7).

##### 3. Testing how individual investment in cooperative care varies with group size

The effect of group composition on the time individuals invested in incubation was modelled using a BLMM with a binomial error distribution. The response variable was the number of observation minutes an individual was observed sitting versus the number of minutes it was not sitting, which accounts for variation in the amount of time individuals were observed. The number of males and females in groups were included as fixed effects and year, enclosure, group and individual identity were entered as random effects. Separate models were run for males and females (R code: M7 & M8). For this analysis only data on primary nests were included as attendance at secondary and tertiary nests was sporadic, and the presence of secondary and tertiary nests did not influence the total amount of time groups protected their nests (Tables S6 & S7).

##### 4. Testing how coordination over reproduction changes with group size

The effect of group composition on the number of interruptions to incubation was modelled using a BLMM with a Poisson error distribution. The response variable was the total number of interruptions observed across all observations divided by the number of hours groups were observed (this was multiplied by 100 and rounded to whole numbers as MCMCglmm requires count data to be whole numbers). The number of males and females in groups were included as fixed effects, and year and enclosure were included as random effects (R code: M9). We removed five enclosure-by-year records where no incubation was observed as this removes the possibility for interruption. The effect of the disparity in the time males and females invested in incubation on the number of interruptions was modelled in the same way, but an extra fixed effect of the difference in the proportion of time males and females spent incubating was included (R code: M10).

##### 5. Testing how coordination over reproduction influences reproductive success

The effect of interruptions on the proportion of eggs broken in nests was modelled using a BLMM with a binomial error distribution. The response variable was the number of eggs broken versus the number of eggs not broken. The difference in the proportion of time males and females spent incubating and the number of males in groups were included as fixed effects, and the year and enclosure were included as random effects (R code: M11). The impact of the broken eggs on the overall hatching success of groups was modelled using a BLMM with a binomial error distribution with the number of eggs hatched versus the number of eggs that did not hatch as the response variable. The proportion of eggs that were broken was included as a fixed effect, and year and enclosure were included as random effects (R code: M12).

##### 6. Supplementary analyses

We present two additional analyses in the supplementary materials (Fig. S1. Table S14 & S15) that are not discussed in the main text, but may provide useful information to some readers. These analyses examine the effects of group composition on the total number of eggs (R code: M13) and chicks (R code: M14) produced by groups as opposed to the per individual measures of reproductive success presented in the main text.

## Data and Code Availability

All data and code are available at the open science framework: DOI 10.17605/OSF.IO/4ZTFN.

## Acknowledgments

We are thankful to Tobias Uller, Philip Downing and Heikki Helanterä for comments on earlier drafts of the manuscript, to all the staff at Oudtshoorn Research Farm for assistance with data collection and maintenance of the birds, and to the Western Cape Government for use of their resources. This research was funded by the Swedish Research Council (grant number 2017-03880), the Knut and Alice Wallenberg Foundation (Wallenberg Academy fellowship numbers 2013.0129 & 2018.0138), Carl Tryggers (grant numbers 12: 92 & 19: 71) to CKC and a stipend from the Carlsberg Foundation to MFS and by the Jörgen Lindström stipendium to JM. The maintenance and development of the ostrich population used in this study was funded by the Western Cape Department of Agriculture and supported by grants from the Western Cape Agricultural Research Trust (Grant number 0070/000VOLSTRUISE: Cloete) as well as the Technology and Human Resources for Industry program (THRIP – Grant number TP14081390585) of the South African National Research Foundation to SWPC.

## Author contributions

Conceptualization JM, SWPC, CKC; Methodology JM, CKC; Formal analysis JM, MFS, CKC; Investigation JM, MFS, MB, SWPC, CKC; Data curation JM, MFS, MB, ZB, AE, SWPC, CKC; Writing original draft JM, CKC; Writing, Reviewing & Editing JM, MFS, MB, ZB, AE, SWPC, CKC; Visualization JM, MFS, CKC; Supervision JM, MFS, CKC; Funding acquisition JM, SWPC, CKC.

